# Keratinocyte Pannexin-1 is essential for Mechanical Hypersensitivity following Traumatic Peripheral Nerve Injury

**DOI:** 10.1101/2025.10.27.684945

**Authors:** Christina M. Mecca, Olena Isaeva, Alexander R. Mikesell, Anvitha Sriram, Bradey A. Stuart, Bhavya Dharanikota, Cheryl L. Stucky

## Abstract

Neuropathic pain remains one of the most prevalent and challenging forms of chronic pain to manage. It is characterized by severe cutaneous touch and cold evoked pain. Whereas the contribution of peripheral nerves to neuropathic pain is well established, the influence of peripheral non-neuronal cells that interact with injured nerves has not been studied in depth. Keratinocytes, the primary cell type in the epidermis, play an important role in the initial processing of environmental mechanical and thermal stimuli under normal conditions. Here, we investigated the role of keratinocytes following traumatic peripheral nerve injury and determined that these cells contribute to injury-associated heightened sensation. We performed tibial spared nerve injury (tSNI) in mice that selectively express archaerhodopsin in keratinocytes to temporally reduce activity from these cells at various timepoints following injury. Brief (5 min) optogenetic inhibition *completely reversed* both the mechanical and cold hypersensitivity at acute (4 days) and throughout chronic (17 weeks) timepoints following tSNI. Keratinocytes isolated from the spared glabrous skin dermatome were sensitized to mechanical and cold stimuli and found to release factors that enhance activity of sensory neurons. Moreover, we found that keratinocyte phospholipase A2 (PLA2) driven activation of pannexin-1 is critical for the long term development of mechanical allodynia. This discovery indicates that keratinocytes contribute substantially to the mechanical and cold allodynia after traumatic nerve injury and suggest that keratinocytes may be a topical therapeutic target to alleviate the severe touch- and cold-evoked cutaneous pain following peripheral nerve injury.

## INTRODUCTION

Each year, traumatic peripheral nerve injuries occur in across the globe[1] and 50-90% of these patients subsequently develop chronic neuropathic pain [2]. Traumatic neuropathic injury produces a range of sensory symptoms, including cutaneous mechanical and cold allodynia, hyperalgesia, dysesthesia, and paresthesia^[3, 4]^.

Current treatments for neuropathic pain—gabapentinoids, tricyclic antidepressants (TCAs), and opioids—are limited by side effects, tolerance, and misuse risk[5]. Thus, it is critical to identify alternative routes to develop more effective treatments. Traumatic neuropathic pain arises from direct nerve insult followed by Wallerian degeneration of injured nerves and formation of neuroma. It is associated with increased neuronal activity within the neuroma, dorsal root ganglia, and spinal cord. Traditionally, cutaneous sensory neurons were believed to be the sole mediators of detecting environmental sensory stimuli, and thus the exclusive mediators of neuropathic pain. However, recent discoveries revealed a crucial role for non-neuronal skin cells in the detection and relay of mechanical and thermal signals to peripheral somatosensory circuit in both healthy and disease states[6–9]. Keratinocytes comprise the majority (90-95%) of epidermal cells, express a variety of sensory receptors, and release a range of neuroactive factors to communicate with sensory afferents and non-neuronal cells in the skin^[7, 10–15]^.

Extracellular application of mechanical and cold stimuli, induce calcium influx, membrane depolarization, and release of neuroactive molecules from keratinocytes^[6, 7, 9, 16]^. Furthermore, keratinocytes contribute to detecting both innocuous and noxious touch and temperature information in healthy skin ^[6, 7, 9]^, but keratinocytes’ roles in heightened pain after tissue injury is early in its exploration.

Emerging evidence suggests a role for keratinocytes in neuropathic pain. Work done by Clark and colleagues[17] assessed the role of human keratinocytes in the context of fracture and CRPS pain, which has a notable skin phenotype in its pathophysiology including changes to epidermal thickness and temperature which is thought to be a result of neurogenic inflammation[18]. Additionally, Wnt5a signaling in keratinocytes as a driver of CRPS-related pain hypersensitivity. Wnt5a upregulated NR2B and MMP9 in both skin and dorsal root ganglia, exacerbating pain.

Inhibition of Wnt5a reduced mechanical and thermal hypersensitivity, implicating keratinocyte-neuron interactions in CRPS pathology [19]. Furthermore, Radtke et al., demonstrated that transplanted human keratinocytes near injured afferents induced extreme neuronal hyperexcitability, spontaneous firing, and pain behaviors in rodent models [20]. Finally, previous work from our lab showed that keratinocytes contribute to chemotherapy-induced peripheral neuropathy (CIPN) through sensitization of PIEZO1, where paclitaxel is given systemically and can directly affect skin cells[21]. It is not known whether keratinocytes contribute to the tactile and cold pain after traumatic nerve injury, where the keratinocytes themselves are not apparently damaged or directly affected by injury. Here we set out to determine whether keratinocytes are sensitized following traumatic nerve injury. We used a spared nerve injury model where two nerves are cut, and degenerate and one nerve remain innervating skin; we focused on the keratinocytes of the skin that remains innervated.

### Animals

Adult male and female C57BL.6J mice aged 8-10 weeks of age were used in the experiments. To selectively inhibit keratinocyte activity, we generated a mouse line expressing Archaerhodopsin-3 (Arch) under the keratin 14 (K14) promoter by crossing Ai35D (B6;129S-Gt(ROSA)26Sor^tm35.1(CAG-aop3/GFP)Hze^/J) Archaerhodopsin (Jackson stock number 012735) and B6N.Cg-Tg(*KRT14*-cre)1Amc (Jackson stock number 018964) mice. Mice were housed in a barrier facility, five animals per cage, at a room temperature of 25 ± 1°C and 25-45% humidity, under a 12/12 h day/night cycle. Animals had *ad libitum* access to food and water, and the cage bedding used was P. J. Murphy Coarse Grad Sani-Chips. Experiments were performed in accordance with the National Institute of Health (NIH) guidelines and approved by the MCW Institutional Animal Care and Use Committee (IACUC) (Milwaukee, WI; Protocol #0383).

### Tibial Spared Nerve Injury model

To induce Tibial Spared Nerve Injury (tSNI) in mice, we used an established protocol [22]. Briefly, the animals were deeply anesthetized with isoflurane (2%) for the duration of the surgery (≤ 15 min). An incision was made on the lateral mid-thigh, and the underlying muscles were separated to expose the three distal branches of the sciatic nerve. The common peroneal and sural nerves were ligated with silk sutures distal to their branch point and transected with 2-3 mm of each nerve excised distal to the ligature. The tibial nerve was preserved and contact with this nerve was avoided. The muscle and skin of the surgical area were closed with sutures after the procedure. For the sham surgery, mice were anesthetized, and their sciatic nerves were exposed but not ligated or cut. All surgeries were performed on the right hind limb of the animals.

### Behavior

For all behavioral assessments, animals were randomly assigned to their treatment groups, with the experimenter blinded to treatment/condition identity until after the behavioral data were collected and analyzed. Behavioral assessments were conducted between 8:00 a.m. and 3:00 p.m. Mice were habituated to the experimental room and testing chambers, either on a mesh wire rack or a glass top surface, for 2 h prior to testing, with the experimenter present for the last 30 min of the acclimation period.

#### von Frey up/down test

Mechanical sensitivity was assessed in the hind paw of mice using the von Frey up/down test, as previously described [23]. Briefly, the hind paws of the mice were stimulated with calibrated von Frey filaments (0.02–1.4 g). A withdrawal response was characterized by an up/down motion of the stimulated hind paw; paw sliding or toe flaring was not considered a response. The 50% withdrawal threshold was determined after data collection.

#### Cold plantar assay

To assess cold sensitivity, we used the dry ice cold plantar assay, according to an established protocol[9]. The mice were tested on a glass top surface (2.5 mm). A syringe (10 mL) containing densely packed powdered dry ice was applied to the glass below the right hind paw. Withdrawal latency was recorded in three trials. The glass top was allowed to warm back to room temperature between trials.

#### Hargreaves assay of radiant heat

The Hargreaves thermal heat assay was used to determine thermal heat sensitivity[9, 24]. Similar to the dry ice test, the mice were placed on a glass top surface. Radiant heat was applied to the right hind paw under glass using an incandescent light bulb. Withdrawal latencies were measured in three trials. The glass top was allowed to warm back to room temperature between trials.

#### Drug delivery

Apyrase, ^10^PanX, and conditioned media were delivered subcutaneously into the mouse hind paw, as previously described[7, 9]. Insulin Syringes were loaded with 20 µL of the injection solution and kept on ice until the time of injection. The experimenter was blinded to the identity of the injection until data collection and analysis. Hind paw injections Apyrase from potato (Sigma A6237) was solubilized, aliquoted, and stored according to the manufacturer’s recommendations. On the day of the experiment, 10 units of apyrase in PBS or vehicle (PBS) were prepared and loaded into syringes. Animals underwent baseline behavior assessment prior to drug injection and were tested 45 min after injection. For cell culture medium injections, keratinocyte cultures from sham or tSNI mice were collected and filtered. Media per animal was aliquoted in mL in Eppendorf tubes, flash frozen, and stored at −80°C until use. Blank unconditioned media controls were prepared in the same manner as the conditioned media groups. Animals were given a pre-injection baseline on the day of experimentation and were tested at 0.5, 1, 2, 3, 24, and 48 hr after injection for mechanical sensitivity; cold sensitivity was tested at 0.5, 1, and 2 hr post-injection. For hind paw injection of ^10^PanX, stock solutions and aliquots were prepared according to the manufacturer’s recommendations. On the day of the experiment, the baseline behavior of the mice was recorded prior to a 20 µL hind paw injection of 100 µM ^10^PanX. Behavior was recorded again one hour after the injection.

### Cell Culture

#### Epidermal keratinocytes

The tibial region of the right hind paw skin of mice was collected and placed in an Eppendorf tube containing 10 mg/mL dispase (Gibco, ThermoFisher Scientific, Waltham, MA, USA) and Hanks Balanced Salt Solution (HBSS) (without CaCl_2_, MgCl_2_, or MgSO_4_) for 45 min. The dermis and epidermis were separated using fine forceps. The epidermis was incubated in a plastic Petri dish containing HBSS (without CaCl_2_, MgCl_2_, or MgSO_4_), 0.05% Trypsin, and 50% EDTA for 27 min at room temperature. Trypsin was neutralized by the addition of 15% heat-inactivated FBS. Keratinocytes were dissociated from the epidermal sheet by mechanical rubbing using a disposable p1000 pipette tip, as previously described [25]. Once dissociated, the HBSS keratinocyte solution was centrifuged for 6 min at 0.3 rcf. The supernatant was discarded, and the cells resuspended in warmed EpiLife medium supplemented with 1% human keratinocyte growth supplement (HKGS), 0.2% Gibco Amphotericin B, and 0.25% penicillin-streptomycin. The cells were then plated on laminin coated coverslips (Calcium imaging and Patch Clamp recordings), or PLL-collagen coated wells and grown at 37°C and 5% CO_2_ until confluent (96-well plate assays).

#### Sensory neurons

Naïve C57 mice were deeply anesthetized under isoflurane and sacrificed for extraction of DRG neurons as previously described. Briefly, DRGS at lumbar levels 1-6 were collected and incubated in collagenase type IV (1 mg/mL) for 40 min in an incubator kept at 37°C and 5% CO2. Next, DRGs were trypsinized in 0.05% trypsin for 45 min. Afterwards trypsin was neutralized with High Horse Serum before undergoing gentle mechanical dissociation. Dissociated neurons were plated onto laminin coated glass coverslips and allowed to seed for 2 hours prior to being fed with complete medium (Dulbecco’s modified Eagle’s medium (DMEM)/Ham’s F12 medium supplemented with 10% heat-inactivated horse serum, 2 mM l-glutamine, 1% glucose, penicillin (100 U/ml), and streptomycin (100 μg/ml)). Cells were allowed to grow overnight in an incubator kept at 37°C and 5% CO2.

#### Sensory neuron – epidermal keratinocyte co culture

For sensory neuron-keratinocyte co-cultures, each cell type was collected as described above. Naïve sensory neurons were plated first and allowed to seed before addition of sham or tSNI keratinocyte cultures. The plated cells were seeded for two hours before being fed with EpiLife medium supplemented with 1% human keratinocyte growth supplement (HKGS), 0.2% Gibco Amphotericin B, and 0.25% penicillin-streptomycin. Co-cultures were allowed to grow over night at 37°C and 5% CO2.

### Calcium imaging and cold ramp stimulation

#### General protocol + Tibial SNI and Sham Keratinocytes

Cells were loaded with the ratiometric calcium dye, FURA-2AM (2.5 µg/mL Fura-2 in 2% BSA) for 45 minutes. The cells were washed for an additional 30 min before imaging. Extracellular buffer containing (in mM) 150 NaCl, 10 HEPES, 8 glucose, 5.6 KCl, 2 CaCl2, and 1 MgCl2 (pH 7.4, 320 Osm) was used for perfusion during experiments (6 mL/min). The extracellular perfusion buffer was equilibrated to room temperature before testing. The room temperature was measured each test day to ensure the starting point matched the environment the coverslips were incubating in (∼21-24 °C). To test calcium responses to cold stimuli, a cold ramp stimulus was created using an in-line cooler– with the ramp starting at room temperature and dropping to ∼12 C° over 3 min. Temperature of buffer was recorded via LabChart software. The calcium responses were recorded using a Nikon Eclipse TE200 inverted microscope with Nikon Elements imaging software (Nikon Instruments, Melville, NY), with each cell showing at least a 30% increase in its 340/380 nm ratio being counted as a calcium response.

### Electrophysiology

#### Keratinocyte Mechanosensitivity Recordings

Keratinocytes were isolated from the glabrous skin of adult C57BL/6J mice on day 4 after tibial spared nerve injury (SNI) or sham surgery, as previously described. Cells were cultured for 24 - 48 hours before recordings. Whole-cell voltage-clamp recordings (holding potential −40 mV) were performed using borosilicate glass pipettes filled with intracellular solution containing (in mM): 135 KCl, 1 MgCl₂, 1 EGTA, 0.2 NaGTP, 2.5 Na₂ATP, 10 glucose, and 10 HEPES (pH 7.25–7.35). Cells were perfused with extracellular HEPES-buffered solution (in mM): 127 NaCl, 3 KCl, 2.5 CaCl₂, 4 MgCl₂, 10 HEPES, and 10 glucose (pH 7.35–7.40). Mechanical stimulation was applied using a piezo-driven glass probe (PA25, PiezoSystem Jena) delivering 150 ms step displacements in 0.125 μm increments at 10 s intervals.

#### Sensory Neuron Excitability Recordings

Whole-cell patch-clamp recordings in current-clamp mode were performed on small-diameter DRG neurons (≤32 μm) co-cultured with tSNI or sham keratinocytes 24 hours after plating. Neurons were superfused with extracellular solution containing (in mM): 140 NaCl, 3 KCl, 2 CaCl₂, 1 MgCl₂, 10 HEPES, and 10 glucose (pH 7.4). Patch electrodes (4–8 MΩ) were pulled from borosilicate glass and filled with internal solution containing (in mM): 135 KCl, 4 MgCl₂, 2 EGTA, 0.2 NaGTP, 2.5 ATPNa₂, and 10 HEPES (pH 7.2). Whole-cell access was first established in voltage-clamp mode, then switched to current-clamp to measure resting membrane potential. Spontaneous action potentials (APs) were recorded for 2 minutes without current injection. Cells firing ≥1 AP during this period were classified as spontaneously active. To assess evoked excitability, cells were held at −70 mV and stimulated with a series of 500-ms depolarizing current steps (0–450 pA in 25-pA increments). All recordings were performed at room temperature using an Axopatch 200B amplifier and pCLAMP software (Molecular Devices).

### Bulk mRNA Sequencing

Mice were deeply anesthetized with isoflurane before cervical dislocation. Glabrous skin innervated by the Tibial nerve was isolated from the right hind paw of ten-week-old male C57 mice that underwent tSNI or Sham surgery and incubated in 10 mg/mL dispase dissolved in HBSS for 45 min to 1 hour at room temperature. The epidermis was separated from the dermis and immediately flash frozen. Samples were kept at −80°C until transport to the Mellows Center at MCW. RNA extraction was performed using the Qiagen RNAeasy Plus Kit following the kits instructions. Paired sequencing was performed using the NovaSeq platform. The generated sequencing data was exported as FastQ files for analysis. Partekflow transcriptomic analysis software was used to analyze the mRNA sequencing data in the FastQ files. Prealignment and trimming of unaligned reads were performed using the Partek recommended parameters.

Alignment of reads was performed using the splice-aware aligner, STAR (2.7.8a). The dataset was then aligned using genome build of mus musculus (mouse) mm39. Post alignment (QA/QC) reports were generated. After alignment, the sequencing data was then quantified to an annotated model (mm39 - RefSeq Transcripts98) using the Partek recommended parameters. Gene counts were normalized as counts per million (CPM) which was the recommended method by Partekflow. Differential expression analysis (DEA) was used to generate a list of differentially expressed genes (DEGs) from the dataset. The DEGs were then uploaded to Ingenuity Pathway Analysis (IPA) for further exploration of biologically relevant pathways.

### PLA2 Activity Assay ELISA

Whole epidermis was collected as previously described in bulk sequencing experiments. After epidermal sheet collection, the samples were processed using bead homogenizer. Samples were plated in triplicate and prepared following the Phospholipase A2 Activity Assay Kit (ab273278, Abcam) instructions. Once samples were pipetted into their assigned wells (tSNI vs sham with and without wasp venom). The plate was read using a Spectramax plate reader in kinetic mode for 50 min. After 50 minutes the data was exported and a standard curve generated. A random point before the curve plateau was selected and used for analysis (20 min timepoint). The relative enzymatic activity was calculated based off of the 20 min time point data according to manufacturer protocol. A Pierce BCA Protein Assay (ThermoFisher) was run on each sample to determine total protein content per sample. The PLA2 plate run data was normalized per sample by their total protein content.

### YO-PRO-1 Dye uptake assay

YO-PRO-1 dye uptake assay (Y3603, ThermoFisher) was modified based on previous studies [26]. Briefly, primary keratinocyte cultures from 5 male naïve C57 mice were performed and plated on a black 96 well plate with PLL and Collogen. Cells were allowed to grow to confluency for 5-7 days. On the day of the assay, keratinocytes were washed three times with Hank’s Balance Salt Solution containing Calcium to remove any cell media. Cells were then allowed to equilibrate for 3 minutes before being treated with 5 µM YO-PRO-1 and their assigned treatment (vehicle, Melittin, ^10^PanX, or combination). Each treatment was performed in triplicate. The treatments were allowed to incubate at room temperature with gentle rocking. After 10 minutes the plate was read at (give wavelengths) for 40 minutes. At 25 minutes Triton-X was added to lyse the cells and increase YO-PRO-1 signal as a positive control. Run data was then graphed and analyzed. Twenty minutes was used to calculate the change in fluorescence compared to the first reading of the run as 20 minutes what the time point used for calculating activity of PLA2. For tSNI vs Sham keratinocytes, the assay was run in basal conditions.

### Cytokine Array

After 24 hours in culture, DRG culture conditioned media was collected, filtered to remove debris, and flash frozen and stored at −80°C until time for experimentation. A mouse sandwich ELISA cytokine array (C3 RayBiotech) was used to profile inflammatory mediators present in the DRG media following the kit instructions. Blots were quantified via ImageJ (Fiji Film).

### qRT-PCR

The glabrous hind paw skin (of B6PX Cre+ and Cre - mice given tSNI or sham surgeries) was dissected and the epidermis was dissociated from the dermis using Dispase (Gibco 17105-041; 20mg/mL). RNA was isolated using TRIzol and a PureLink RNA Mini Kit (Invitrogen, Life Technologies). Total RNA content was assessed with a Nanodrop Light Spectrophotometer (Thermo Scientific). cDNA was synthesized using the Superscript VILO cDNA Synthesis Kit (Invitrogen, Life Technologies). The quantitative real-time PCR reaction was performed using a Bio-Rad CFX384 Touch Real-Time PCR Detection System. All samples were run in triplicate and gene expression was analyzed using the ΔΔCt method and normalized to GAPDH. The primers used were obtained from Integrated DNA Tehcnologies: *mPannexin*1-qF: AAGCAGATCCAGTCCAAG; *mPannexin1*-qR: GGGCTCTTCTCCTTCTCC; *mGAPDH*-qF: AGGTCGGTGTGAACGGATTTG; *mGAPDH*-qR: TGTAGACCATGTAGTTGAGGTCA.

### Data Analysis

All statistical analyses were performed using GraphPad Prism (GraphPad Software, La Jolla, CA). Data were assessed for normality prior to statistical testing, and appropriate parametric or non-parametric tests were applied. Statistical significance was defined as p < 0.05. Results are presented as mean ± SEM, unless otherwise indicated.

#### Behavioral Testing

Mechanical sensitivity assessed via the von Frey up/down method was analyzed using two-way repeated-measures ANOVA with Sidak’s post hoc test. Media behavior assays were evaluated using two-way repeated-measures ANOVA with Tukey’s post hoc test. Behavioral responses following apyrase injection and optogenetic stimulation were analyzed using three-way repeated-measures ANOVA, with Tukey’s post hoc test applied for optogenetic data. Calcium Imaging.

#### Electrophysiology and Imaging

Skin-nerve recordings were analyzed using two-way repeated-measures ANOVA with Sidak’s post hoc test for repeated measures, and unpaired Student’s t-tests for threshold comparisons. Patch clamp mechanical thresholds and current amplitudes were analyzed using Mann–Whitney U-tests. Proportional responses to mechanical stimuli and current profiles were compared using Chi-squared tests with Fisher’s Exact post hoc correction. Mechanically activated (MA) currents were compared using unpaired Student’s t-tests. Co-culture recordings were analyzed using Fisher’s Exact test for spontaneous activity, Welch’s t-test for action potential counts, Student’s t-test for resting membrane potential, and two-way repeated-measures ANOVA for input-output curves. Yoda-1 calcium imaging data were analyzed using one-way ANOVA with Bonferroni post hoc test for group comparisons, and Chi-squared tests with Fisher’s Exact correction for proportional data. Cold calcium imaging and YO-PRO-1 uptake assays were analyzed using unpaired Student’s t-tests or two-way repeated-measures ANOVA, depending on the experimental design. Naïve keratinocyte–DRG media calcium imaging was analyzed using one-way repeated-measures ANOVA.

#### Molecular Analysis

Messenger RNA expression levels were compared using unpaired Student’s t-tests. Cytokine array data were analyzed by calculating fold changes between sham and tSNI conditions, with a 2-fold change threshold used to define biologically relevant alterations.

## RESULTS

### Inhibition of epidermal keratinocytes completely reverses long-lasting peripheral nerve injury-induced mechanical and cold hypersensitivity

Since patients who endure traumatic nerve injury typically experience mechanical allodynia, we first asked whether keratinocyte activity is required for evoked mechanical hypersensitivity after traumatic peripheral nerve injury. To selectively inhibit keratinocyte activity, we used a transgenic mouse line expressing archaerhodopsin-3 (Arch) driven by a keratin-14 (K14) promoter.

Archaerhodopsin is a light-sensitive proton pump that, when activated, causes cell membrane hyperpolarization via hydrogen ion efflux [34]. This hyperpolarization inhibits depolarization-dependent cellular activity, effectively inhibiting keratinocyte signaling[1]. Arch-expressing mice (Arch-K14Cre^+^) and littermate controls (Arch-K14Cre^-^) were subjected to tibial spared nerve injury (tSNI), a preclinical model that induces robust mechanical and cold hypersensitivity, modeling a Grade V nerve injury in humans[2] ^[3, 4]^. Starting one week after surgery, we used the up-down von Frey method to assess mechanical sensitivity prior to and during keratinocyte inhibition. To transiently inhibit the activity of keratinocytes, amber light was applied for six minutes before and during stimulation. We found that Arch-K14Cre+ mice at the pre-surgery baseline showed a significant increase in mechanical and cold withdrawal thresholds, similar to what we have shown previously [11–12]. Furthermore, sham control Arch-K14Cre+ mice also showed the same finding throughout the experimental time course (Fig. 1 A, D). Brief inhibition of keratinocyte signaling *fully* reversed nerve injury-induced mechanical hypersensitivity from early (week 1) through chronic (week 17) timepoints; withdrawal thresholds of Arch-K14Cre-controls were unaffected (Fig. 1A).

**Figure 1.**
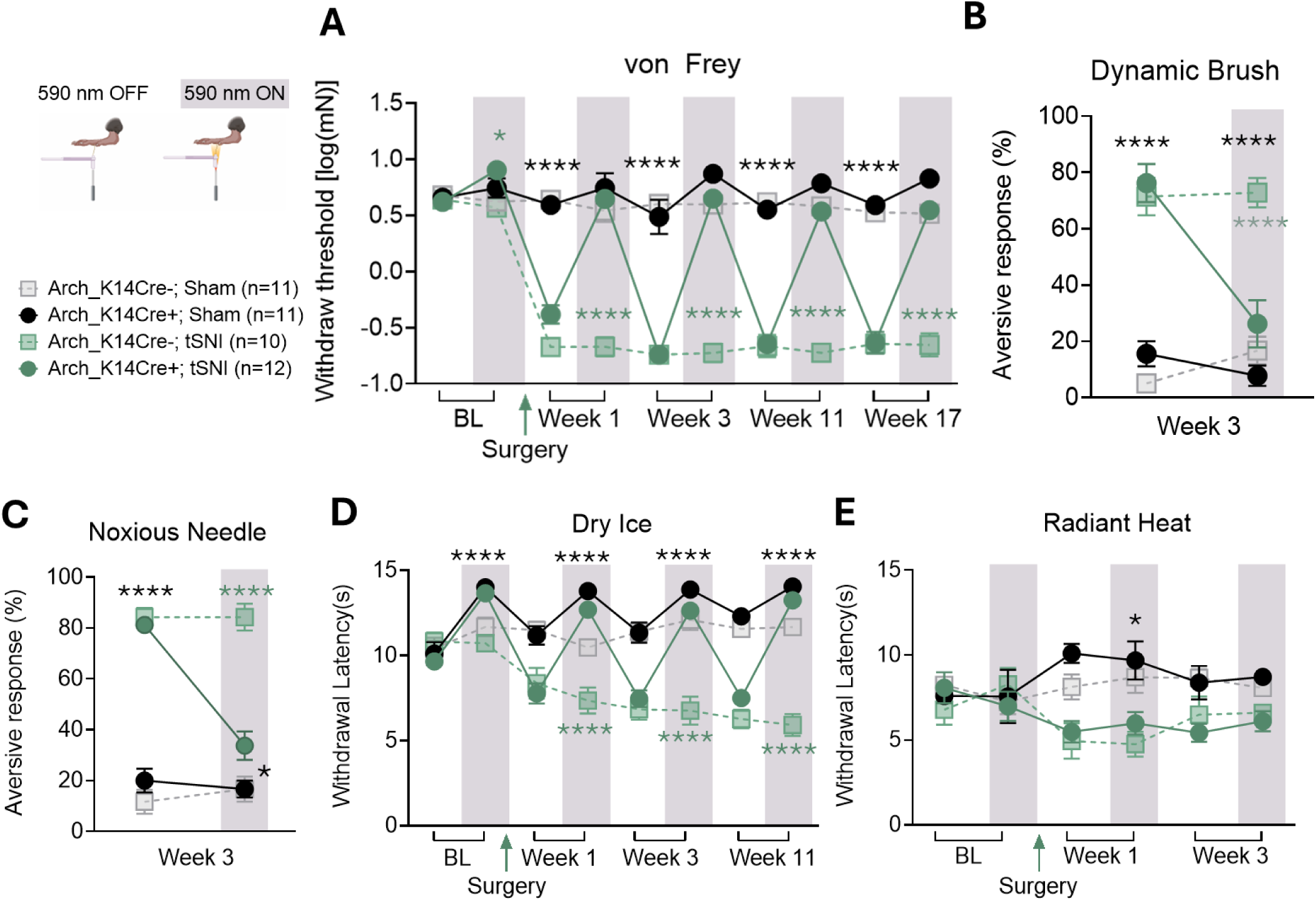
Optogenetic inhibition of keratinocyte activity fully reverses mechanical and cold hypersensitivity but does not affect thermal hypersensitivity following peripheral nerve injury. **(A)** Summary data of mechanical withdrawal thresholds recorded from sham and tSNI Arch-K14Cre- and Arch-K14Cre+ mice at baseline (BL) and at weeks 1, 3, 11, and 17 following surgeries. (**B** and **C**) Percentage of aversive attending responses (out of 10 trials) to **(B)** a paintbrush (dynamic light touch) and **(C)** a 25-gauge needle (noxious needle test). Recordings were made in the presence or absence of amber light stimulation at Week 3 following surgery. Sample size: n= 10-12. **(D)** Cold withdrawal latency from the dry ice plantar assay at BL and 1-, 3-, and 11-weeks post-surgery, collected in the presence or absence of hind paw stimulation of the mouse hind paws. Sample size: n=10-12 for (A,D) n=7-8 for (B,C). **(E)** Withdrawal latency to radiant heat taken at BL, 1-, and 3-weeks post-surgery. All data were collected with and without amber light stimulation of the mouse hind paw to inhibit keratinocyte activity. 590 nm OFF = clear, 590 nm ON = grey; sample size: for (E) n=7-8 (male and female mice). Stats: 3-Way RM ANOVA with Tukey’s post-hoc *p < 0.05, **p < 0.01, and ****p < 0.0001 for A-E, for **(A)** Testing Point p<0.0001, Injury p<0.0001, Genotype p<0.0001, Testing Point x Injury p<0.0001, Testing Point x Genotype p<0.0001, Injury x Genotype p<0.0001, Testing Point x Injury x Genotype p<0.0001 **(B)** Testing Point p=0.019, Injury p<0.0001, Genotype p=0.035, Testing Point x Injury p=0.0014, Testing Point x Genotype p=0.0004, Injury x Genotype p=0.0063, Testing Point x Injury x Phenotype p=0.04,**(C)** Testing Point p=0.0008, Injury p<0.0001, Genotype p=0.0009, Testing Point x Injury p=0.0003, Testing Point x Genotype, Injury x Genotype p<0.0001, Testing Point x Injury x Phenotype p=0.004 **(D)** Testing Point p<0.0001, Injury p<0.0001, Genotype p<0.0001, Testing Point x Injury p<0.0001, Testing Point x Genotype p<0.0001, Injury x Genotype p<0.0001, Testing Point x Injury x Phenotype p=0.0004 **(E)** Testing Point p=0.95, Injury p<0.0001, Genotype p=0.43, Testing Point x Injury p=0.0004, Testing Point x Genotype p=0.43, Injury x Genotype p=0.51, Testing Point x Injury x Genotype p=0.61. All data in this figure are represented as mean ± SEM, unless otherwise stated.

To test other forms of evoked mechanical stimulation, we also assessed dynamic light brush and noxious pin prick after tibial SNI. The tSNI-enhanced dynamic light brush and pin prick hypersensitivity both nearly completely reversed during keratinocyte inhibition (Fig. 1B and C). These data indicate that keratinocyte activity is essential for the persistent hypersensitivity to punctate mechanical stimuli, dynamic light brush and noxious pin prick. Next, we asked whether keratinocyte activity is essential for the thermal hypersensitivity following nerve injury. Cold hypersensitivity is common in patients with nerve injury^[2, 3, 5, 27]^. To test cold hypersensitivity, mutant mice (Arch-K14Cre+) and their littermate controls (Arch-K14Cre-) were placed on a glass surface, and dry ice was applied under the glass to cool the glabrous hind paw skin. In the absence of amber light, Arch-K14Cre+ mice exhibited cold hypersensitivity within the first week after tSNI. Optogenetic inhibition of keratinocytes *completely* reversed cold hypersensitivity from the first week through the 11^th^ week post-injury (Fig. 1D). In contrast, tSNI-induced heat hypersensitivity was unaffected by amber light exposure (Fig. 1E). These data indicate that ongoing keratinocyte activity is critical for detecting and transducing both evoked mechanical and cold hypersensitivity associated with peripheral nerve injury. In contrast, ongoing keratinocyte activity is dispensable for tSNI-induced heat hypersensitivity or normal heat sensitivity.

### Tibial SNI induces sensitization of keratinocytes to mechanical and cold stimuli

Given that keratinocyte inhibition reverses mechanical and cold hypersensitivity following tSNI, we hypothesized that tSNI-induced hypersensitivity may result, at least in part, from enhanced keratinocyte responsiveness to mechanical and cold stimuli. To test this hypothesis, we isolated keratinocytes from sham and tSNI mice and measured their responses to mechanical and cold stimuli. Keratinocytes were isolated and cultured from the plantar skin region innervated by the spared tibial nerve of tSNI and sham C57BL/6J mice on postoperative day 4 (POD4). To measure mechanical sensitivity, we performed whole-cell patch-clamp recordings on keratinocytes and mechanically stimulated their cellular membranes using a blunt glass probe. Each keratinocyte was tested with stepwise increasing membrane indentation, and the amount of membrane indentation required to elicit an inward current was recorded. If a keratinocyte did not respond to the highest indentation level, it was considered mechanically insensitive (Fig. Fig. 2A) shows an example of mechanically activated (MA) currents induced in keratinocytes in response to a stepwise increase in membrane indentation. Our data indicate that the proportion of keratinocytes responding to mechanical stimulation was significantly higher in the tSNI group than in the sham controls (Fig. 2B). No differences in MA current density or mechanical threshold were observed between the treatment groups (Fig. 2C and D). To determine whether tSNI affects keratinocyte sensitivity to cold, we used calcium imaging to measure keratinocyte responsiveness to a ramp of cooled buffer (from 22-24°C to 11-12 C° over 3 min). All keratinocytes from tSNI or sham animals responded to the cooled buffer with an increase in intracellular calcium concentration (Fig. 2E). Summary data indicated that keratinocytes from tSNI mice exhibited responses to milder temperatures (indicating increased sensitivity to cold) compared to those from sham mice (Fig. 2F). However, the magnitude of the calcium response did not differ between the treatment groups (Fig. 2G). Together, these data indicate that keratinocytes from tSNI animals are intrinsically sensitized to both mechanical and cold stimuli in the skin.

**Figure. 2.**
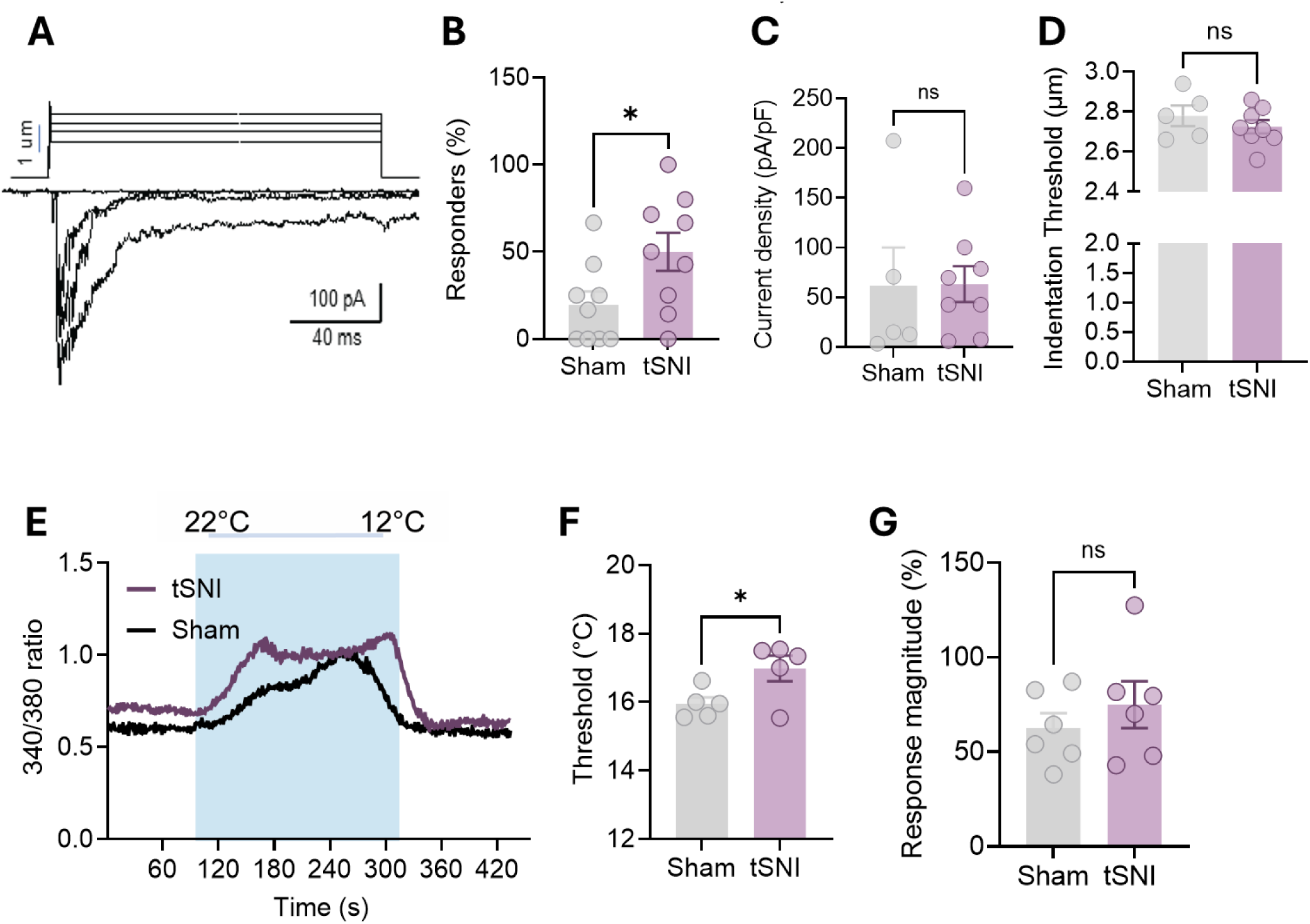
Tibial SNI induces keratinocyte sensitization to mechanical and cold stimuli. **(A)** Protocol for whole-cell patch-clamp recording of mechanically activated (MA) currents evoked by increasing membrane indentation in keratinocytes isolated from a sham mouse on postoperative day 4 (POD4). **(B)** Percentage of keratinocytes that responded to membrane indentation with MA current. Circles depict the average count of responsive keratinocytes per mouse **(B, C, D)**. **(C)** Comparative analysis of maximal MA current density **(D)** and activation threshold recorded from keratinocytes isolated from sham and tSNI mice on POD2-POD4. Circles represent the average MA current density and threshold per animal. Sample size: n = 5-8 mice per group, 3 to 8 cells per animal. **(E)** Example of calcium responses to cold ramp stimulation recorded from keratinocytes isolated from sham and tSNI mice on POD4. **(F, G)** Comparative analysis of threshold **(F)** and magnitude **(G)** of keratinocytes’ calcium responses to cold stimulation. Circles depict the average thresholds and magnitudes of calcium responses observed in keratinocytes for each animal. Sample size: n = 5 mice per group, 60 to 90 keratinocytes per animal. Unpaired Student’s t-test, for **(B)** p=0.37, **(C)** p=0.04, **(D)** p=0.97, **(F)** p=0.04, **(G)** p=0.42. All data in this figure are represented as mean ± SEM, unless otherwise stated.

### Tibial SNI keratinocytes release factors that enhance naïve neuronal excitability and influence mechanical behavior

Since keratinocytes exhibited mechanical and cold sensitization, we asked whether crosstalk between tSNI keratinocytes and sensory neurons is altered. To test this, we isolated keratinocytes from the hind paw of either tSNI or sham mice and co-cultured them with lumbar dorsal root ganglia (DRG) sensory neurons derived from naïve mice. Twenty-four hours after plating, whole-cell patch-clamp recordings were performed on the neurons (Fig. 3A). We found that a higher proportion of DRG neurons co-cultured with tSNI keratinocytes exhibited spontaneous action potential (AP) firing compared to those co-cultured with sham keratinocytes; spontaneous activity frequency did not differ between treatment groups (Fig. 3B, C, D).

**Figure 3.**
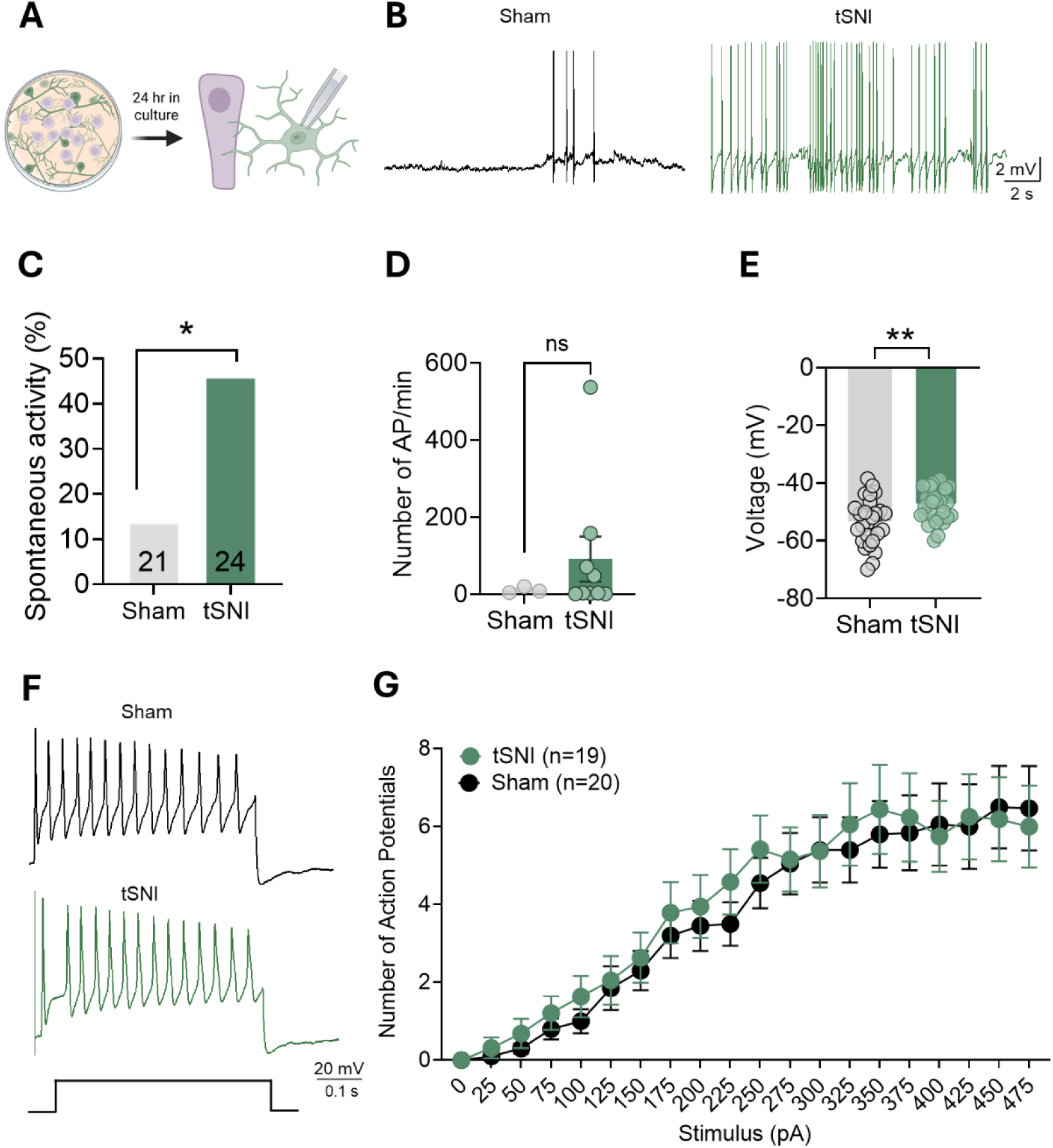
Tibial SNI keratinocytes release factors that enhance excitability of naïve sensory neurons. **(A)** Experimental schematic. **(B)** Representative traces of spontaneous action potential firing recorded from DRG neurons co-cultured with tSNI and sham keratinocytes**. (C)** A greater proportion of DRG neurons co-cultured with tSNI keratinocytes overnight exhibited spontaneous action potential activity. **(D)** Number of action potentials fired per minute. **(E)** Depolarized resting membrane potential**. (F)** Representative traces of action potential firing in response to a 500 ms step, 450 pA current injection, recorded from DRG neurons co-cultured with tSNI and sham keratinocytes**. (G)** Averaged input-output curves plotting the number of action potentials in response to stepwise current injections. Sample size: n = 4 mice per group; n = 4-6 recordings per mouse. *p < 0.05, **p < 0.01. Statistics: **(C)** Fisher’s exact test; p=0.03 **(D)** Welch t-test p=0.2041. **(E)** Student’s t-test; p=0.003. **(G)** 2-Way RM ANOVA, Stimulus p<0.0001, Injury p=0.54, Stimulus x Injury p=0.99. All data are represented as mean ± SEM.

Additionally, sensory neurons co-cultured with tSNI keratinocytes exhibited more depolarized resting membrane potentials than those co-cultured with sham keratinocytes (Fig. 3E). To evaluate current-evoked firing frequency of sensory neurons, we held the membrane potential at −70 mV and applied a series of 500 ms current injections ranging from 25 pA to 475 pA in 25 pA increments (Fig. 3F). Summary data indicate that sensory neurons co-cultured with tSNI and sham keratinocytes showed no differences in the AP firing frequencies in response to depolarizing steps (Fig. 3G). These data indicate that short-term co-culturing of naïve DRG neurons with keratinocytes from tSNI mice leads to membrane depolarization and increased spontaneous firing of sensory neurons, indicating enhanced basal excitability. However, the capacity of these tSNI keratinocyte-cultured neurons to generate APs in response to depolarizing inputs remains unaltered.

Next, we asked whether factors released by keratinocytes from tSNI mice induce mechanical and cold hypersensitivity. To test this, keratinocytes from the glabrous hind paw of C57BL/6 mice with tSNI or sham surgery were isolated on POD4 and cultured *in vitro*. Conditioned media from tSNI or sham keratinocytes were collected on day 3 post-plating, filtered, and injected into the hind paw of naïve C57BL/6J mice. Von Frey mechanical and plantar cold assays were conducted before and after injection of tSNI or sham conditioned, or blank media. We determined that the blank and sham control injected groups did not have any significant differences in mechanical or cold responses, thus we proceeded to make comparisons between the sham and tSNI injected groups (Fig S1). Our data indicate that injection of tSNI-conditioned media into the hind paw of naïve mice was sufficient in inducing mechanical hypersensitivity (Fig 4A), whereas cold sensitivity remained unchanged (Fig. S1B). These data suggest that factors released by tSNI keratinocytes can sensitize naïve mice to mechanical stimuli.

**Figure. 4.**
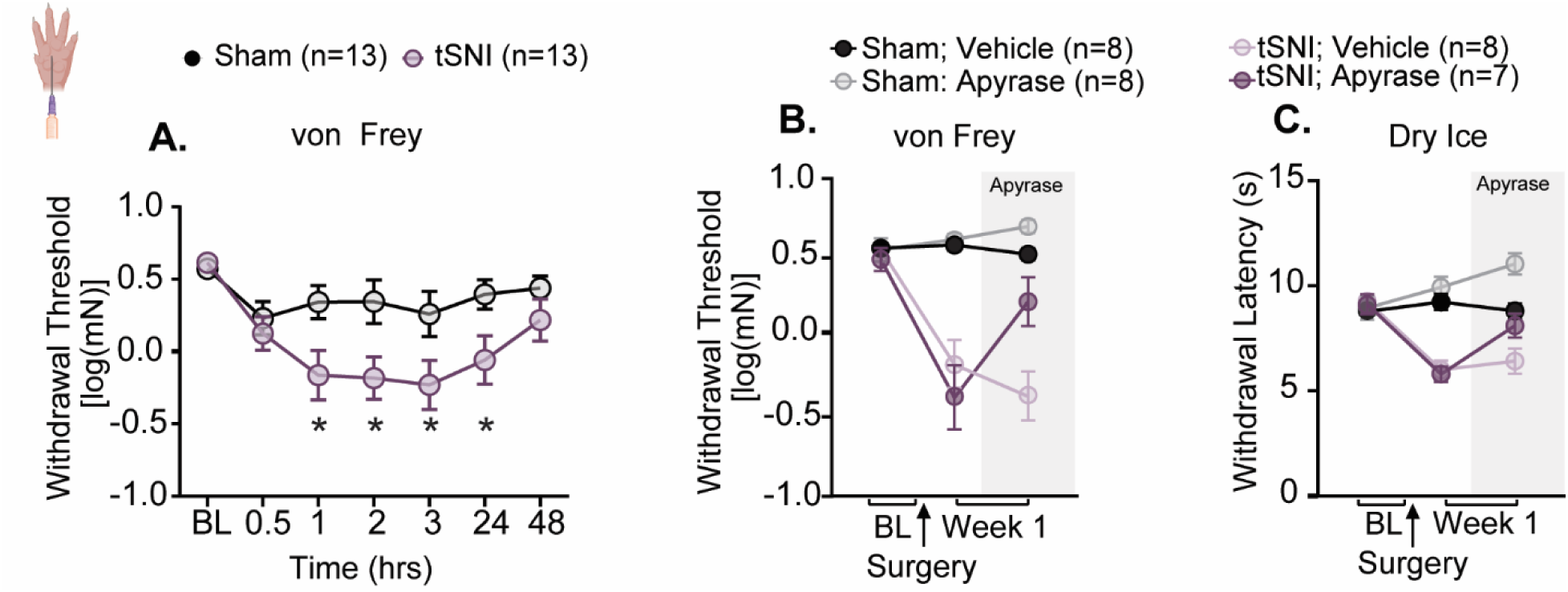
tSNI keratinocytes release factors that induce mechanical hypersensitivity that appear to be independ of epidermal ATP. Naïve C57BL/6 mice were assessed for **(A)** mechanical sensitivity (von Frey test). Behavioral response before and after hind paw injection of 10 units of apyrase or vehicle for **(B)** mechanical (von Frey) and **(C)** cold (cold plantar assay) at 1-week post-surgery. Statistics: For **(A)** Two-way repeated-measures ANOVA was performed using Tukey’s post-hoc test. Time × Treatment p=0.022, Time p<0.0001, Treatment p=0.019. For **(B),** three-way repeated-measures ANOVA Time × Treatment × Injury p =0.87, Time × Treatment p=0.002, Injury × Treatment p=0.39, Time × Injury p<0.0001, Treatment p=0.01, Injury p=<0.0001, Time p=0.002. For **(C)** Three-way repeated measure ANOVA Time × Treatment × Injury p =0.84, Time × Treatment p=0.03, Injury × Treatment p=0.19, Time × Injury p<0.0001, Treatment p=0.02, Injury p<0.0001, Time p=0.005. All data in this figure are represented as mean ± SEM, unless otherwise stated..

### ATP signaling does not play an additional role in mechanical and cold responses after nerve injury

We previously demonstrated that mechanical activation of keratinocytes triggers the release of ATP, which in turn activates P2X4 receptors on sensory neurons of naïve, uninjured animals[7]. Thus, ATP is a candidate transmitter for paracrine signaling between keratinocytes and sensory afferent endings following nerve injury. Since ATP plays a role in normal keratinocyte responses to mechanical and thermal stimuli^[7, 9]^, we investigated the extent to which ATP signaling contributes to mechanical hypersensitivity following tSNI. Within the first week post-surgery, von Frey mechanical and cold assays were conducted on tSNI or sham mice before and after intraplantar injection of apyrase, an enzyme that hydrolyzes extracellular ATP to ADP and eventually to adenosine. Apyrase treatment did not significantly alter the mechanical withdrawal thresholds (Time × Treatment × Injury p =0.87) (Fig. 4B) or increase the paw withdrawal latency to cold stimulation (time × treatment × injury p =0.84) (Fig. 4C). Therefore, while ATP signaling is important for normal mechanical and thermal detection of stimuli as previously described[11–12], local degradation of ATP does not appear to play a critical role in tSNI induced hypersensitivity.

### PLA2 activity is enhanced after tibial SNI

To better understand how factors that are potentially released from keratinocytes following nerve injury, we performed bulk mRNA sequencing of the ipsilateral hind paw epidermis isolated from tSNI and sham mice on postoperative day 4 (POD4) (Fig. 5). Bulk sequencing analysis revealed various differentially expressed genes (DEGs) (Fig. 5A). These genes were then used for pathway analysis using the Ingenuity Pathway Analysis (IPA) software. Phospholipase A2 (PLA2) was identified as both an upstream regulator and part of a causal network based on genes that were differentially expressed after tSNI (Krtap6-2, Krtap6-1, Krtap6-5, and Spprb2) (Fig. 5B). These regulators and networks predict the activity/involvement of molecules that may behave differently despite not showing changes in transcript expression. PLA2 activity leads to increased production of arachidonic acid and lysophospholipids – both pathways have been shown to be involved in pain signaling[28]. To validate the predicted involvement of PLA2, we performed a PLA2 activity assay on the epidermis collected from tSNI and sham mice (Fig. 5C). Indeed, PLA2 activity was increased in the tSNI samples (Fig. 5C).

**Figure. 5.**
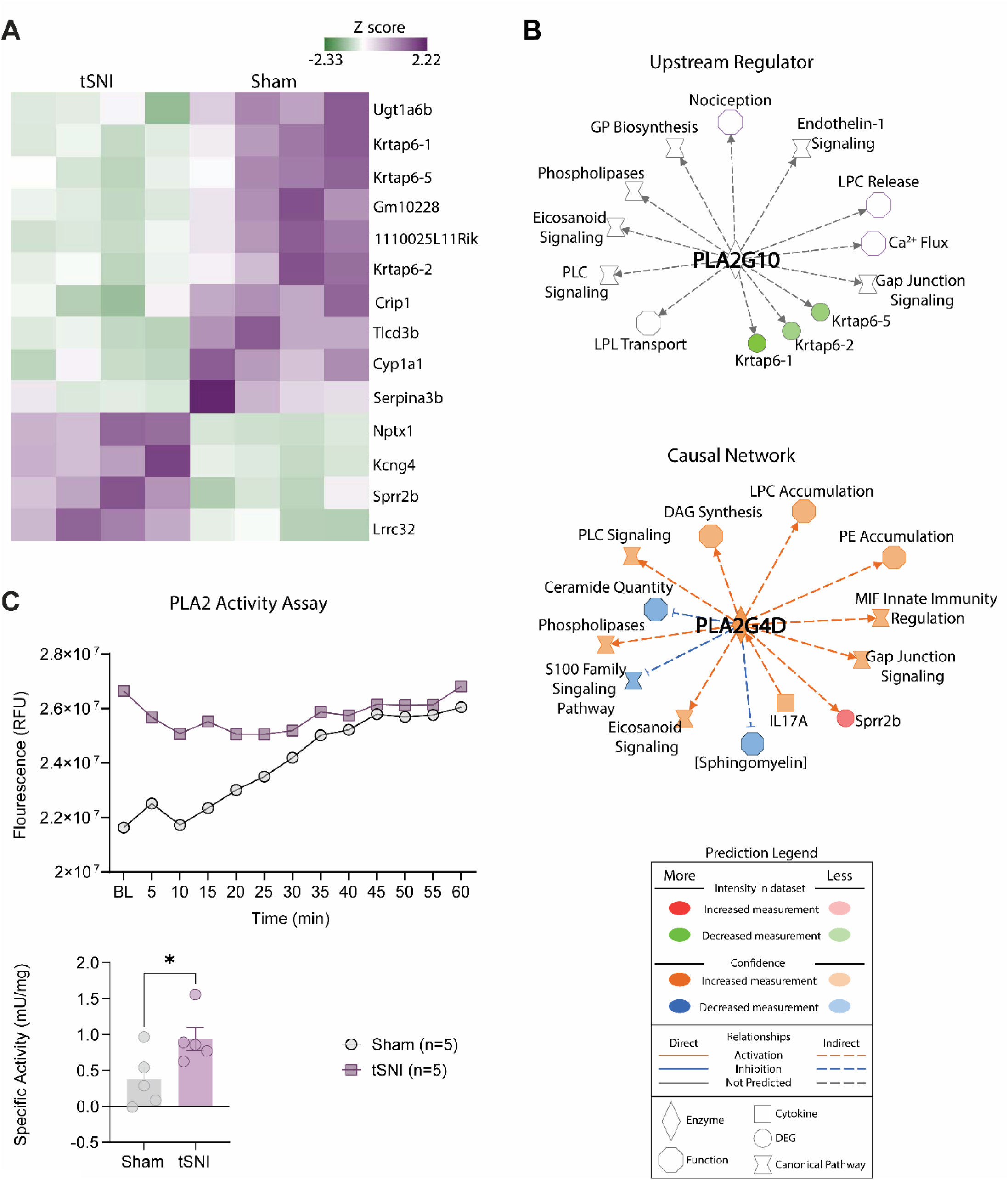
PLA2 is both an upstream regulator and part of the upstream causal network that may contribute to keratinocyte sensitization. Epidermal samples from tSNI and sham mice on postoperative day 4 (POD4) were used for transcriptomic analysis **(A)** Heatmap of differentially expressed genes (DEGs) genes are represented as Z-scores using the normalized genes in counts per million (CPM) **(B)** Pathway map generated in Ingenuity Pathway Analysis. Differentially expressed genes Krtap6-2 (p = 1.29E-05), Krtap6-1 (p = 2.29E-08), and Krtap6-5 (p = 0.0000099) predicted PLA2 as an upstream regulator (PLA2G10, p = 1.72E-05) (top) and as part of a causal network (PLA2G4D, p = 1.55E-02) (bottom). Abbreviations: GP, glycerophospholipid; LPC, lysophosphatidlycholine; LPL = Lipoprotein lipase; MIF = Macrophage Migration Inhibitory Factor, PE, phosphatidylethanolamine; DAG, diacylglycerol; sprr2b, small proline-rich protein 2B. For casual networks: lysophosphatidlycholine p= 8.88E-05, ceramide p=1.61E-05, phosphatidylethanolamine p=0.000266, sphingomyelin p=0.000266, diacylglycerol p= 0.00337, sprr2b p = 3.97E-08, **(C)** Functional validation of PLA2 in tSNI and Sham isolated epidermal samples. The specific activity was calculated using plate read data 20 min after the start of the assay. Two-tailed unpaired t-test, p=0.0444; sample size: n=5 mice per group. *p < 0.05, **p < 0.01, and ****p < 0.0001. All data in this figure are represented as mean ± SEM, unless otherwise stated.

### PLA2 is nessicary and sufficienct in inducing Pannexin 1 activation in keratinocytes

PLA2 is involved in inflammation and pain signaling[9,10]; therefore, we investigated whether PLA2 activity could influence the release of neuroactive factors after tSNI. Interestingly, the accumulation of lysophospholipid lysophosphatidylcholine (LPC) was predicted based on the predicted activation of cytosolic (c)PLA2, Pla2G4D. Recently, Henze et al. (2025) showed that PLA2 driven production of lysophospholipids, particularly LPC, can open pannexin channels. Pannexin-1 is a large pore-forming channel that releases a variety of neuroactive mediators, including ATP, upon activation [12]. This channel is highly expressed in keratinocytes, plays an important role in their mechanosensitivity, and has been implicated in pain signaling[29–34]. To determine whether PLA2 driven activation of pannexin-1 occurs in keratinocytes, we performed a YO-PRO-1 dye uptake assay as previously described[26] (Fig. 6A-C). YO-PRO-1 is a fluorescent dye that can pass through large pore-forming channels, such as pannexin channels, to interact with the cell’s DNA, producing a fluorescent signal. If pannexin-1 is opened, an increase in the YO-PRO-1 signal is detected. Keratinocytes isolated from the hind paw of naïve C57BL/6 mice were cultured and treated with either vehicle or melittin, a potent PLA2 activator, in the presence or absence of the pannexin-1 inhibitor ^10^PanX. We observed a significant increase in YO-PRO-1 dye uptake with 5 µM melittin, which was completely prevented when pretreated with ^10^PanX, confirming that YO-PRO-1 signal generation was produced by pannexin-1 channel opening. We also utilized a concentration of melittin (1µM) that was below the activation threshold for PLA2[22] and observed no changes to YO-PRO-1. Next, we evaluated whether tSNI keratinocytes had more open pannexin-1 channels compared to sham. We saw that tSNI keratinocytes indeed have increased YO-PRO-1 signal at rest, suggesting more pannexin-1 channels are opened (Fig 6D-E). Together, these findings support our hypothesis that PLA2 activity leads to pannexin-1 activation in keratinocytes and that pannexin-1 is more active after tSNI.

**Figure. 6.**
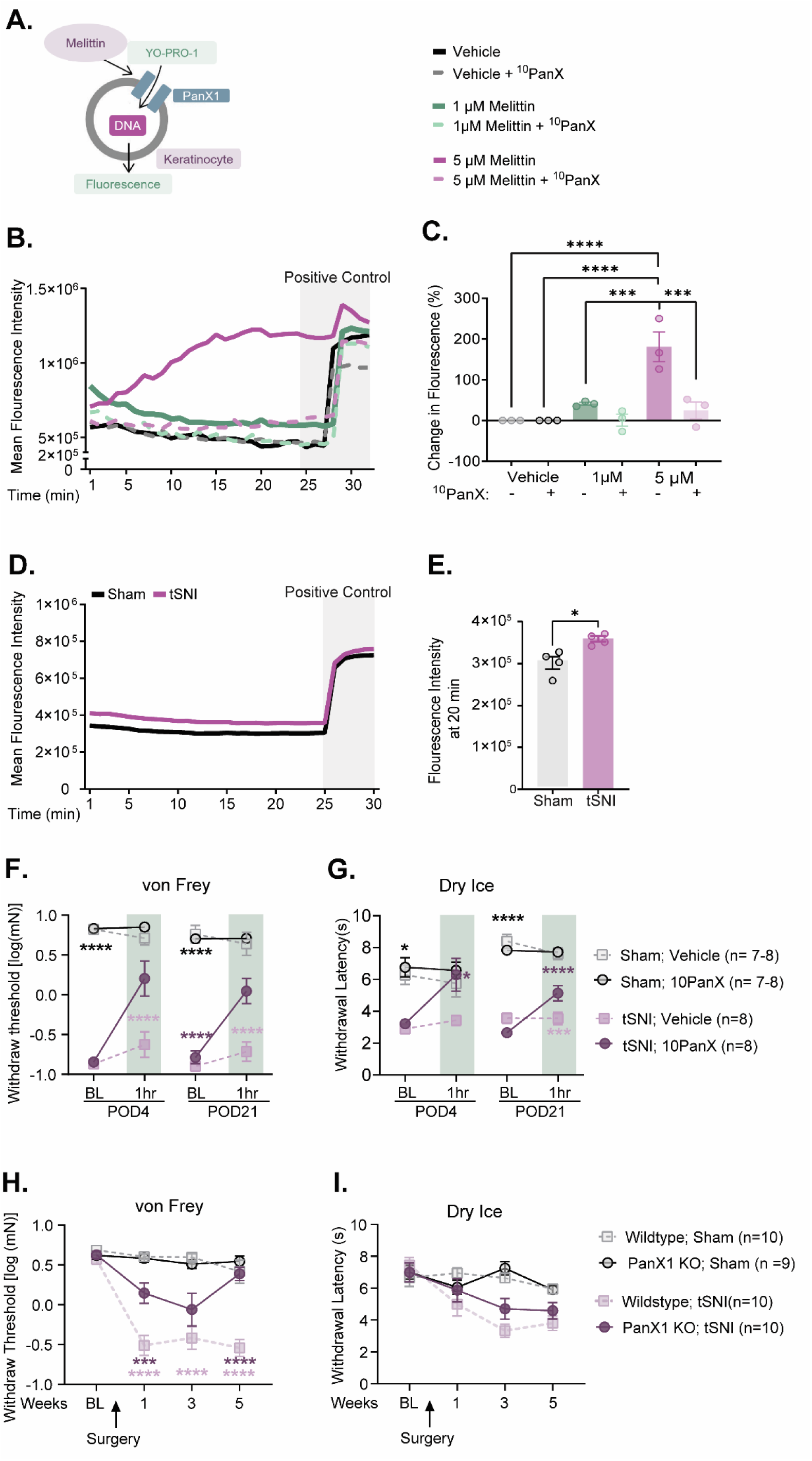
Keratinocyte pannexin-1 is important for injury-induced *mechanical* hypersensitivity. YO-PRO-1 dye uptake assay. **(A)** When pannexin-1 channels are opened, YO-PRO-1 is able to pass through the cell and interact with DNA to generate a fluorescent signal. YOPRO-1 dye uptake responses in naïve mouse keratinocytes treated with vehicle, melittin with and without ^10^PanX. **(B)** Time course of YOPRO-1 dye uptake in naïve mouse keratinocytes treated with vehicle and melittin with and without pannexin-1 inhibition **(C)** Percent change in fluorescence signal (*e.g.* pannexin-1 channel opening) compared to that in vehicle-treated group. 2-way RM ANOVA, ^10^PanX × melittin concentration p=0.005, ^10^Panx p=0.002, melittin concentration p= 0.006; sample size; n = 3 per group, males used. **(D)** YOPRO-1 dye uptake responses in tSNI or sham isolated keratinocytes. **(E)** Fluorescence intensity of tSNI vs sham keratinocytes at 20 minutes. Unpaired t-test p= 0.015, n = 4 per group, both sexes used. C57 mice with tSNI or sham surgery before and after 100 µM ^10^PanX hind paw injection **(F)** Mechanical withdrawal thresholds and **(G)** cold withdrawal latencies. Behavioral assessment of mice with epidermal deletion of pannexin-1 with tSNI or sham surgery **(H)** von Frey and **(I)** cold plantar assay. Behavioral testing was conducted before and at weeks 1, 3, and 5 post-surgery 3-way RM ANOVA, for **(H,I) (H)** Time p=0.0003, Injury p<0.0001, Injection p=0.0002, Time × Injury p<0.0001, Time × Injection p=0.007, Injury × Injection p=0.0001, Time × Injury × Injection p=0.023, for **(I)** Time p=0.0003, Injury p<0.0001, Injection p=0.018, Time × Injury p<0.0001, Time × Injection p<0.0001, Injury × Injection p=0.1, Time × Injury × Injection p=0.015. Sample size: n = 7-8 per group, equal sexes used Three-way RM ANOVA. For **(E-F), (E)** Time × Injury × Genotype p= 0.055, Injury × Genotype p <0.0001, Time × Injury p <0.0001, Time p<0.0001, Injury p<0.0001, Genotype p <0.000. For **(F)** Time × Injury × Genotype p=0.3410, Injury × Genotype p=0.19, Time × Genotype p=0.0002, Time × Injury p=0.44, Genotype p <0.0001, Injury p=0.2077, Time p <0.0001; sample size: n = 8 per group, equal sexes used. *p < 0.05, **p < 0.01, and ****p < 0.0001. Pink * = Injury comparison, Purple * = Injury x Genotype All data in this figure are represented as mean ± SEM, unless otherwise stated.

### Keratinocyte pannexin-1 is essential for peripheral nerve-induced mechanical hypersensitivity

Next, we asked whether epidermal pannexin-1 contributes to tSNI-induced mechanical and cold hypersensitivity. To address this, we assessed the mechanical and cold sensitivity of tSNI or sham mice before and after intraplantar injection of 200 µM ^10^PanX. Peripheral inhibition of pannexin-1 significantly attenuated both mechanical and cold responses in tSNI mice on both POD 4 and 21 (Fig. 6 C-D). In contrast, inhibition of pannexin-1 did not affect the behavior of sham mice, a finding consistent with previous studies that showed a minimal role for pannexin-1 in uninjured conditions^[31, 35, 36]^. These data suggest that peripheral pannexin-1 is important for mediating injury-induced mechanical and cold hypersensitivity. Since hind paw injection of the pannexin-1 inhibitor could act on sensory terminals and other cells residing in the epidermis, we generated a keratinocyte-specific knockout of pannexin-1 using the K14 Cre driver (PanX1-K14 Cre+ or −) to determine whether, and to what extent, keratinocyte pannexin-1 is involved in injury-induced hypersensitivity. The deletion of pannexin-1 from keratinocytes was validated using qRT (Fig. S2). The PanX1-K14 Cre+ and littermate control (PanX1-K14 Cre-) mice underwent tSNI or sham surgery, and their mechanical and cold sensitivity were assessed before and during 1-, 3-, and 5-weeks post-surgery. Behavioral assessment revealed that keratinocyte deletion of pannexin-1 significantly alleviated mechanical hypersensitivity during the first three weeks post-injury (Fig. 6E). Moreover, PanX1_K14 Cre+ mice showed full recovery of normal mechanosensitivity by 5 weeks post-tSNI. These findings suggest that keratinocyte pannexin-1 plays a critical role in the development and maintenance of nerve injury-induced mechanical allodynia. In contrast, deletion of pannexin-1 in keratinocytes had no effect on the cold hypersensitivity that developed after tSNI (Fig. 6F) or did it affect baseline responses to mechanical or cold stimuli (Fig. 6). These findings provide mechanistic insights into the distinct cellular and molecular pathways underlying nerve injury-induced mechanical and cold hypersensitivity, highlighting pannexin-1’s involvement in mechanical pain.

## DISCUSSION

In summary, the present study shows keratinocytes are critical for both mechanical and cold hypersensitivity induced by traumatic nerve injury. Furthermore, PLA2-Pannexin-1 signaling is critical for mechanically evoked pain developed after traumatic nerve injury. Deletion of pannexin-1 from keratinocytes is sufficient in preventing the development of long-term mechanically evoked hypersensitivity. Additionally, activation of PLA2 in keratinocytes induces YO-PRO-1 dye uptake in a pannexin-1 dependent manner. Thus, we have identified a novel mechanism in which mechanically evoked pain is generated and maintained via a keratinocyte PLA2-Pannexin-1 signaling axis. This novel discovery is the first of its kind to show keratinocyte involvement after traumatic nerve injury, not only providing a mechanistic explanation for keratinocyte involvement, but also, identifying a potential topical therapeutic target to address the severe cutaneous touch evoked pain after traumatic nerve injury. This suggests that keratinocyte signaling is not only necessary for the initiation but also for the maintenance of pain behaviors.

Our work further explores the role of non-neuronal cell involvement in neuropathic pain, addressing the question of skin involvement in pain after traumatic nerve injury.

Keratinocytes are known to detect mechanical, thermal, and chemical stimuli and communicate with sensory afferents via mediators such as ATP^[7, 9]^. Previous work from our lab demonstrated that optogenetic inhibition of keratinocytes blunts mechanically evoked membrane depolarization and ATP release. Building on this, our current findings show that keratinocyte hyperpolarization via archaerhodopsin leads to full reversal of mechanical and cold hypersensitivity, indicating a direct role of keratinocyte signaling in evoked hypersensitivity.

Interestingly, keratinocytes became sensitized to mechanical and cold stimuli following tibial spared nerve injury (tSNI), despite remaining uninjured. This suggests that the injury-induced environment, including factors released from degenerating and intact nerves, may reprogram keratinocytes, leading to altered gene expression and heightened sensitivity to mechanical and cold stimuli. These changes could amplify their role in sensory detection and contribute to pain hypersensitivity.

We further demonstrated that keratinocytes from tSNI mice release factors that enhance the intrinsic excitability of naïve sensory neurons. Co-culture experiments revealed increased spontaneous action potential firing and depolarized resting membrane potentials in neurons exposed to tSNI keratinocytes. This supports the hypothesis that keratinocytes can modulate neuronal activity through paracrine signaling, potentially contributing to peripheral sensitization. Furthermore, conditioned media from tSNI keratinocytes induced mechanical hypersensitivity in naïve mice, although cold sensitivity remained unaffected. This suggests that soluble keratinocyte-derived factors are sufficient to induce mechanical hypersensitivity. The lack of cold hypersensitivity may be due to several factors, including rapid degradation of cold-relevant mediators, insufficient concentration to reach activation thresholds, or the requirement for cell– cell contact or spatially restricted signaling that is lost in conditioned media injection experiments. Additionally, cold hypersensitivity may involve distinct molecular pathways or receptor populations that are not activated by the soluble factors present in the media, warranting further investigation into keratinocyte signaling in cold pain.

While ATP is a well-established mediator of keratinocyte-to-neuron communication, our data indicate that ATP degradation via apyrase does not significantly alter mechanical or cold hypersensitivity after tSNI. This implies that ATP may contribute to baseline sensitivity but is not the sole driver of injury-induced hypersensitivity. Bulk mRNA sequencing of the epidermis revealed PLA2 as an upstream regulator, and PLA2 activity was confirmed to be elevated post-injury.

We showed that PLA2 activation leads to pannexin-1 channel opening in keratinocytes, as evidenced by YO-PRO-1 dye uptake assays. This activation was blocked by the pannexin-1 inhibitor ^10^PanX, confirming the specificity of the response. These findings suggest that PLA2 activity sensitizes pannexin-1 channels and contribute to keratinocyte-mediated pain signaling. Finally, we demonstrated that keratinocyte-specific deletion of pannexin-1 significantly reduced mechanical hypersensitivity and prevented its long-term development after tSNI. These results highlight pannexin-1 as a key mediator of keratinocyte-driven pain signaling. Since cold hypersensitivity was unaffected by pannexin-1 deletion, this points to distinct molecular pathways underlying mechanical and cold pain, with pannexin-1 specifically involved in mechanical hypersensitivity.

In conclusion, our findings reinforce the concept that keratinocytes are active contributors to peripheral and potentially central sensitization in neuropathic pain. The identification of a PLA2– pannexin-1 signaling axis provides a mechanistic framework for understanding keratinocyte involvement in traumatic nerve injury and opens new avenues for developing targeted peripheral analgesics.

Overall, our findings demonstrate that keratinocytes, despite remaining structurally intact after nerve injury, undergo sensitization that alters sensory neuron excitability and drives mechanical and cold hypersensitivity. Furthermore, we identify a mechanism in which mechanical hypersensitivity is generated and maintained via PLA2–pannexin-1 signaling. These findings establish keratinocytes as key players in traumatic neuropathic pain and highlight their potential as peripheral targets for analgesic strategies.

## Acknowledgements

The authors would like to thank Stucky Lab members Dianise M. Rodríguez García, Meghan Konda, Samuel J. Zorn, Vanessa Ehlers, and Jonathan Enders for assistance in blinding operators to injury/genotype/intervention prior to experimentation and until after data analysis was completed for behavioral and imaging experiments. The authors would like to thank Anthony Menzel and Vivien Blecking for assistance in cell culture and genotyping. Dianise M. Rodríguez García, Jonathan Enders, and Katelyn Sadler who all participated in or consulted in discussions about data collection and analysis. Members of the MCW Mellows Center for RNA extraction and generation of the initial Meta data to generate FASTQ files for the bulk mRNA sequencing experiment. Jonathan Enders and Michele Battel provided guidance on proper analysis of Bulk mRNA sequencing data. Aniko Szabo of the MCW Biostatistics department provided council on proper statistical analysis for behavioral experiments. MCW Engineering Lab for technical support and generation of fiberoptic lead cables for optogenetic experiments, whose work is supported in part by the NIH shared equipment grant 1S10OD032136-01 3D Printer and the Advancing a Healthier Wisconsin Endowment Project Improving Heart Health, Supporting Healthy Minds & Dismantling Cancer.

## Figures

**Supplemental Fig. 1.**
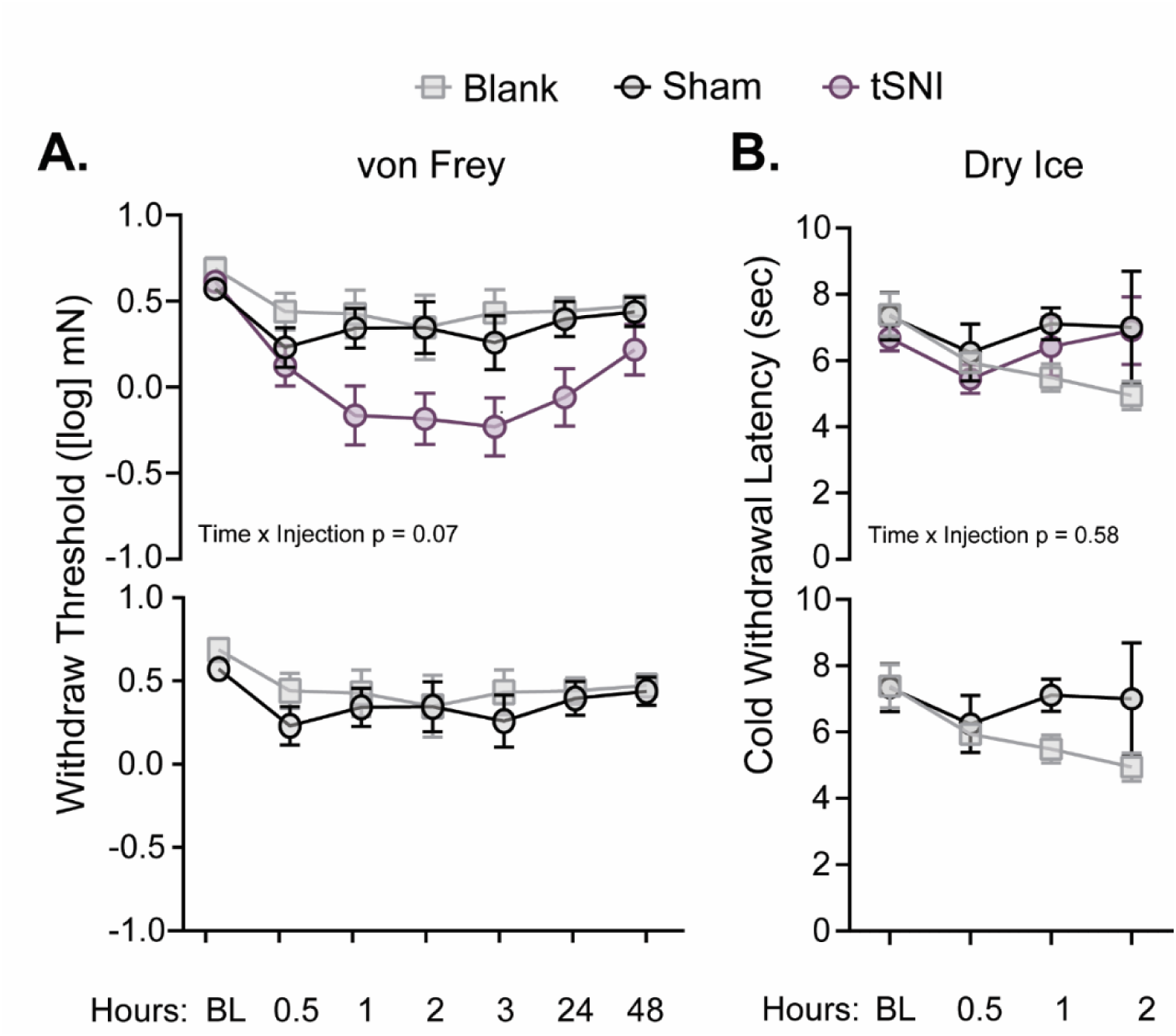
tSNI keratinocytes release factors that induce mechanical but not cold hypersensitivity in naïve mice. Naïve C57BL/6 mice were assessed for **(A)** mechanical sensitivity (von Frey test) and **(B)** cold sensitivity (cold plantar assay) at baseline and after hind paw injection of 20 µL filtered conditioned tSNI or sham or blank control filtered cell culture media; sample size: media: n=3, animals: n=8-13 for **(A)** and n=5 for **(B).** Two-way repeated-measures ANOVA was performed using Tukey’s post-hoc test. For **(A)** Time × Treatment p=0.07, Time p<0.0001, Treatment p=0.01(Top), Time × Treatment p=0.95, Time p=0.1, Treatment p=0.41 (bottom). **(B)** Time × Treatment p=0.52, Time p=0.22, Treatment p=0.37 (Top). Time × Treatment p= 0.44, Time p=0.26, Treatment p=0.19 (bottom). All data in this figure are represented as mean ± SEM, unless otherwise stated.

**Supplemental Fig 2.**
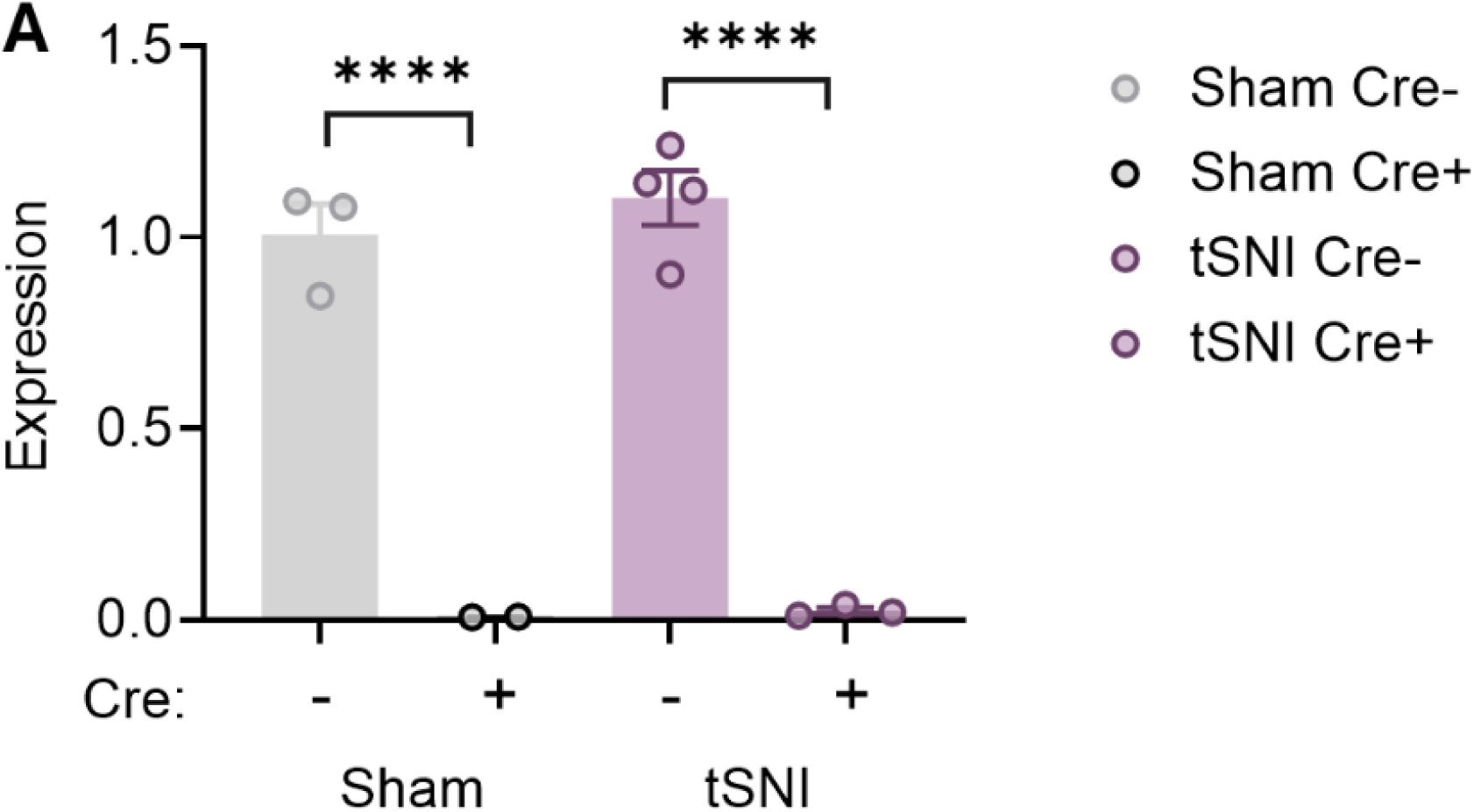
Pannexin-1 is deleted from keratinocytes. Validation of pannexin-1 knock out from keratinocytes. **(A)** Expression of Pannexin-1 in epidermal samples collected from PX1_K14Cre+ or PanX1_K14Cre- mice that received tSNI or sham surgery was performed via mRNA extraction and qRT-PCR. Two-way repeated measures ANOVA was performed with Tukey’s Post Hoc test, Genotype x Injury p=0.56, Injury p=0.43, Genotype p<0.0001; sample size: n=2-4 animals per group

## References

[1] D.J.O. Matos Cruz AJ, Neurotmesis, in: StatPearls (Ed.), StatPearls Publishing, StatPearls [Internet]. 2023.

[2] P.E. Doneddu, U. Pensato, A. Iorfida, C. Alberti, E. Nobile-Orazio, A. Fabbri, A. Voza, Neuropathic Pain in the Emergency Setting: Diagnosis and Management, Journal of Clinical Medicine 12(18) (2023) 6028.

[3] J.N. Campbell, R.A. Meyer, Mechanisms of neuropathic pain, Neuron 52(1) (2006) 77–92.

[4] R.M. Menorca, T.S. Fussell, J.C. Elfar, Nerve physiology: mechanisms of injury and recovery, Hand Clin 29(3) (2013) 317–30.

[5] E. Cavalli, S. Mammana, F. Nicoletti, P. Bramanti, E. Mazzon, The neuropathic pain: An overview of the current treatment and future therapeutic approaches, Int J Immunopathol Pharmacol 33 (2019) 2058738419838383.

[6] K.M. Baumbauer, J.J. DeBerry, P.C. Adelman, R.H. Miller, J. Hachisuka, K.H. Lee, S.E. Ross, H.R. Koerber, B.M. Davis, K.M. Albers, Keratinocytes can modulate and directly initiate nociceptive responses, Elife 4 (2015) e09674.

[7] F. Moehring, A.M. Cowie, A.D. Menzel, A.D. Weyer, M. Grzybowski, T. Arzua, A.M. Geurts, O. Palygin, C.L. Stucky, Keratinocytes mediate innocuous and noxious touch via ATP-P2X4 signaling, Elife 7 (2018) e31684.

[8] Z. Pang, T. Sakamoto, V. Tiwari, Y.S. Kim, F. Yang, X. Dong, A.D. Güler, Y. Guan, M.J. Caterina, Selective keratinocyte stimulation is sufficient to evoke nociception in mice, Pain 156(4) (2015) 656–665.

[9] K.E. Sadler, F. Moehring, C.L. Stucky, Keratinocytes contribute to normal cold and heat sensation, Elife 9 (2020) e58625.

[10] N. Cirillo, The Local Neuropeptide System of Keratinocytes, Biomedicines 9(12) (2021).

[11] C. Erbacher, S. Britz, P. Dinkel, T. Klein, M. Sauer, C. Stigloher, N. Üçeyler, Interaction of human keratinocytes and nerve fiber terminals at the neuro-cutaneous unit, bioRxiv (2022) 2022.02.23.481592.

[12] D. Roggenkamp, S. Köpnick, F. Stäb, H. Wenck, M. Schmelz, G. Neufang, Epidermal nerve fibers modulate keratinocyte growth via neuropeptide signaling in an innervated skin model, J Invest Dermatol 133(6) (2013) 1620–8.

[13] A.T. Slominski, R.M. Slominski, C. Raman, J.Y. Chen, M. Athar, C. Elmets, Neuroendocrine signaling in the skin with a special focus on the epidermal neuropeptides, American Journal of Physiology-Cell Physiology 323(6) (2022) C1757–C1776.

[14] M. Talagas, N. Lebonvallet, R. Leschiera, P. Elies, P. Marcorelles, L. Misery, Intra-epidermal nerve endings progress within keratinocyte cytoplasmic tunnels in normal human skin, Experimental Dermatology 29(4) (2020) 387–392.

[15] M. Talagas, N. Lebonvallet, R. Leschiera, G. Sinquin, P. Elies, M. Haftek, J.P. Pennec, D. Ressnikoff, V. La Padula, R. Le Garrec, K. L’Herondelle, O. Mignen, L. Le Pottier, N. Kerfant, A. Reux, P. Marcorelles, L. Misery, Keratinocytes Communicate with Sensory Neurons via Synaptic-like Contacts, Ann Neurol 88(6) (2020) 1205–1219.

[16] M. Denda, S. Nakanishi, Do epidermal keratinocytes have sensory and information processing systems?, Experimental Dermatology 31(4) (2022) 459–474.

[17] W.W. Li, T.Z. Guo, X.Q. Li, W.S. Kingery, D.J. Clark, Fracture induces keratinocyte activation, proliferation, and expression of pro-nociceptive inflammatory mediators, Pain 151(3) (2010) 843–852.

[18] D.A. Guthmiller KB, Dey S, et al., Complex Regional Pain Syndrome., Treasure Island (FL): StatPearls Publishing, StatPearls [Internet], [Updated 2025 May 4].

[19] H. Zhu, B. Wen, J. Xu, L. Xu, Y. Huang, Wnt5a in keratinocytes contributes to complex regional pain syndrome through the activation of NR2B and MMP9 in rats, Reg Anesth Pain Med (2025).

[20] C. Radtke, P.M. Vogt, M. Devor, J.D. Kocsis, Keratinocytes acting on injured afferents induce extreme neuronal hyperexcitability and chronic pain, PAIN 148(1) (2010) 94–102.

[21] A.R. Mikesell, E. Isaeva, M.L. Schulte, A.D. Menzel, A. Sriram, M.M. Prahl, S.M. Shin, K.E. Sadler, H. Yu, C.L. Stucky, Increased keratinocyte activity and PIEZO1 signaling contribute to paclitaxel-induced mechanical hypersensitivity, Science Translational Medicine 16(777) (2024) eadn5629.

[22] V.L. Ehlers, A. Sriram, B.A.R. Stuart, C.M. Mecca, C.L. Stucky, Sensory neuron PIEZO1 deletion inhibits dynamic light touch sensitivity in uninjured mice, prevents neuropathic light touch hypersensitivity, and drives compensatory changes in dorsal root ganglia, Pain (2025).

[23] K.E. Sadler, F. Moehring, S.I. Shiers, L.J. Laskowski, A.R. Mikesell, Z.R. Plautz, A.N. Brezinski, C.M. Mecca, G. Dussor, T.J. Price, J.D. McCorvy, C.L. Stucky, Transient receptor potential canonical 5 mediates inflammatory mechanical and spontaneous pain in mice, Science Translational Medicine 13(595) (2021) eabd7702.

[24] K. Hargreaves, R. Dubner, F. Brown, C. Flores, J. Joris, A new and sensitive method for measuring thermal nociception in cutaneous hyperalgesia, Pain 32(1) (1988) 77–88.

[25] A.R. Mikesell, O. Isaeva, F. Moehring, K.E. Sadler, A.D. Menzel, C.L. Stucky, Keratinocyte PIEZO1 modulates cutaneous mechanosensation, Elife 11 (2022) e65987.

[26] E. Henze, R.N. Burkhardt, B.W. Fox, T.J. Schwertfeger, E. Gelsleichter, K. Michalski, L. Kramer, M. Lenfest, J.M. Boesch, H. Lin, F.C. Schroeder, T. Kawate, ATP-release pannexin channels are gated by lysophospholipids, bioRxiv (2025) 2023.10.23.563601.

[27] F. Guida, D. De Gregorio, E. Palazzo, F. Ricciardi, S. Boccella, C. Belardo, M. Iannotta, R. Infantino, F. Formato, I. Marabese, L. Luongo, V. de Novellis, S. Maione, Behavioral, Biochemical and Electrophysiological Changes in Spared Nerve Injury Model of Neuropathic Pain, Int J Mol Sci 21(9) (2020) 3396.

[28] J.V. Bonventre, Phospholipase A2 and signal transduction, J Am Soc Nephrol 3(2) (1992) 128–50.

[29] C.Y. Fan, B.B. McAllister, S. Stokes-Heck, E.K. Harding, A. Pereira de Vasconcelos, L.K. Mah, L.V. Lima, N.J. van den Hoogen, S.F. Rosen, B. Ham, Z. Zhang, H. Liu, F.J. Zemp, R. Burkhard, M.B. Geuking, D.J. Mahoney, G.W. Zamponi, J.S. Mogil, S.S. Ousman, T. Trang, Divergent sex-specific pannexin-1 mechanisms in microglia and T cells underlie neuropathic pain, Neuron (2025).

[30] M. Mousseau, N.E. Burma, K.Y. Lee, H. Leduc-Pessah, C.H.T. Kwok, A.R. Reid, M. O’Brien, B. Sagalajev, J.A. Stratton, N. Patrick, P.L. Stemkowski, J. Biernaskie, G.W. Zamponi, P. Salo, J.J. McDougall, S.A. Prescott, J.R. Matyas, T. Trang, Microglial pannexin-1 channel activation is a spinal determinant of joint pain, Science Advances 4(8) (2018) eaas9846.

[31] K.E. Navis, C.Y. Fan, T. Trang, R.J. Thompson, D.J. Derksen, Pannexin 1 Channels as a Therapeutic Target: Structure, Inhibition, and Outlook, ACS Chemical Neuroscience 11(15) (2020) 2163–2172.

[32] S. Penuela, J.J. Kelly, J.M. Churko, K.J. Barr, A.C. Berger, D.W. Laird, Panx1 Regulates Cellular Properties of Keratinocytes and Dermal Fibroblasts in Skin Development and Wound Healing, Journal of Investigative Dermatology 134(7) (2014) 2026–2035.

[33] Q. Wang, H.-y. Li, Z.-m. Ling, G. Chen, Z.-Y. Wei, Inhibition of Schwann cell pannexin 1 attenuates neuropathic pain through the suppression of inflammatory responses, Journal of Neuroinflammation 19(1) (2022) 244.

[34] J.L. Weaver, S. Arandjelovic, G. Brown, S. K. Mendu, M. S. Schappe, M.W. Buckley, Y.-H. Chiu, S. Shu, J.K. Kim, J. Chung, J. Krupa, V. Jevtovic-Todorovic, B.N. Desai, K.S. Ravichandran, D.A. Bayliss, Hematopoietic pannexin 1 function is critical for neuropathic pain, Scientific Reports 7(1) (2017) 42550.

[35] S. Penuela, R. Gehi, D.W. Laird, The biochemistry and function of pannexin channels, Biochimica et Biophysica Acta (BBA) - Biomembranes 1828(1) (2013) 15–22.

[36] Z. Ruan, I.J. Orozco, J. Du, W. Lü, Structures of human pannexin 1 reveal ion pathways and mechanism of gating, Nature 584(7822) (2020) 646–651.

[37] N. Gilbert, F. Chatelain, S. Gibaud, T. Lorivel, Y.-L. Shen, M.-E. Kerros, S. Feliciangeli, F. Fiore, C.-C. Chen, F. Lesage, D. Bichet, Two pore domain THIK2 channel is involved in acute and chronic pain signal regulation, bioRxiv (2025) 2025.08.08.669258.

[38] X.Y. Li, H. Toyoda, Role of leak potassium channels in pain signaling, Brain Res Bull 119(Pt A) (2015) 73–9.

[39] D.V. Vasylyev, P. Zhao, B.R. Schulman, S.G. Waxman, Interplay of Nav1.8 and Nav1.7 channels drives neuronal hyperexcitability in neuropathic pain, J Gen Physiol 156(11) (2024).

[40] J. Sousa-Valente, A.P. Andreou, L. Urban, I. Nagy, Transient receptor potential ion channels in primary sensory neurons as targets for novel analgesics, British Journal of Pharmacology 171(10) (2014) 2508–2527.

[41] Y. Xiao, C. Barbosa, Z. Pei, W. Xie, J.A. Strong, J.M. Zhang, T.R. Cummins, Increased Resurgent Sodium Currents in Nav1.8 Contribute to Nociceptive Sensory Neuron Hyperexcitability Associated with Peripheral Neuropathies, J Neurosci 39(8) (2019) 1539–1550.

[42] S. Baeza-Loya, R.A. Eatock, Effects of transient, persistent, and resurgent sodium currents on excitability and spike regularity in vestibular ganglion neurons, Front Neurol 15 (2024) 1471118.

[43] T. Ngodup, T. Irie, S. Elkins, L.O. Trussell, The Na(+) leak channel NALCN controls spontaneous activity and mediates synaptic modulation by α2-adrenergic receptors in auditory neurons, bioRxiv (2023).

